# Strigolactone receptor DAD2 promotes lateral root formation in *Petunia hybrida*

**DOI:** 10.1101/2024.08.28.609930

**Authors:** Khopeno Khuvung, Pierre-Emmanuel Courty, Didier Reinhardt

## Abstract

Strigolactone regulates various aspects of plant development and interactions with beneficial microbes. While the role of SL in shoot development is highly conserved between monocots and dicots, its role in root system architecture is controversial and context-dependent, although most of the SL appears to be produced there. A central parameter in plant fitness is root-to-shoot ratio, which is determined by growth and branching activity of both, the root as well as the shoot. We used previously characterized mutants defective in SL biosynthesis and sensing to test the role of SL in root growth and branching. Here, we show that a mutant defective in the petunia strigolactone receptor (*dad2*) exhibits defects in root meristem activity and root branching. Notably, this phenotype is not observed in the SL biosynthetic mutant *dad1*, suggesting that DAD2 may perceive a strigolactone analogue that is produced in a DAD1-independently fashion. DAD2 is induced in cortex cells adjacent to lateral root primordia, where it allows them to penetrate the cortex and to grow out of the main root. We propose a model by which inducible SL perception in cells adjacent to LRPs promotes auxin-induced cell wall remodeling in the cortex to allow passage of lateral roots.

**1-sentence summary:** Lateral roots are initiated inside the main root, hence, they have to penetrate several cell layers to grow out. We propose that strigolactone cooperates with auxin to mediate lateral root emergence.

## Introduction

Plant development is under hormonal control at all stages from embryogenesis to organ senescence and programmed cell death, including morphogenesis and regulation of general body plan of the shoot and the root system (Vanstraelen and Benková 2012). A central aspect in this context is branching, which generates characteristic patterns in shoot architecture (Conn et al. 2017; Reinhardt and Kuhlemeier 2002) and in root system organization (Del Bianco and Kepinski 2018; Morris et al. 2017).

A central feature in shoot development is apical dominance, a concept that describes the tendency of the main shoot apex to inhibit the outgrowth of axillary meristems along the stem, thereby favoring the production of a single tall shoot with few or no lateral branches (Domagalska and Leyser 2011; McSteen and Leyser 2005; Rameau et al. 2015). In highly branched shoot systems, apical dominance is decreased, allowing axillary meristems to grow out, and to produce secondary lateral branches in the leaf axils. In the extreme case, this results in recurrent branching at successive hierarchical levels of shoot architecture, giving rise to an exponentially growing number of new shoot tips. This diverts the available resources from the apical meristem, resulting in stunted overall growth of the original main stem. Therefore, highly branched plants are usually dwarfed, which can result in a considerable fitness deficit, although growth under harsh conditions can favor small stature and high branching (Khuvung et al. 2022).

Most plants are somewhere in between, often with a high degree of apical dominance during vegetative development (allowing the plant to compete for light with adjacent neighbors), and a lower degree of apical dominance at the onset of flowering, resulting in increased inflorescence branching and the outgrowth of additional branches from the main stem (Khuvung et al. 2022).

Apical dominance is regulated by several phytohormones, mainly auxin, strigolactone (SL), and cytokinin (CK), but additional hormonal and nutritional factors are involved as well (Beveridge and Kyozuka 2010; Khuvung et al. 2022; McSteen and Leyser 2005; Rameau et al. 2015). Auxin has been identified as the main inhibitory factor that represses the outgrowth of axillary buds along its way of transport by polar auxin transport (PAT) from the site of its biosynthesis in the shoot tip towards the roots. SL is a second inhibitory factor that acts both, directly on axillary buds, and indirectly, by regulation of PAT (Beveridge and Kyozuka 2010; McSteen and Leyser 2005; Rameau et al. 2015). In contrast, CK is a positive regulator of axillary bud outgrowth that acts antagonistically to auxin and SL (Beveridge and Kyozuka 2010; McSteen and Leyser 2005; Rameau et al. 2015).

Root branching is inherently different from shoot branching in several ways: i.) Branching patterns are not determined by the position of organs (the leaves in the shoot), but they are determined by an oscillatory mechanism acting behind the root meristem (Moreno-Risueno et al. 2010); ii.) Lateral root formation requires dedifferentiation of pericycle cells that generate new root meristems (Banda et al. 2019; Du and Scheres 2018), while shoot branching is regulated at the level of outgrowth of preformed axillary meristems; iii.) auxin is an inducer of lateral root formation as opposed to its inhibitory role in bud outgrowth in the shoot (Banda et al. 2019; Du and Scheres 2018).

For SL, a role in root growth regulation has been documented, but a mechanistic understanding of its role in root branching is still lacking. Also, effects of SL vary depending on the plant species studied (Kapulnik et al. 2011; Ruyter-Spira et al. 2011), indicating that the role of SL in root development may be less conserved than in shoot development.

A central role in root development is played by nutrients in the surrounding medium (soil). This represents an important feedback onto root system architecture, which has to be optimized for mineral nutrient and water absorption (Giehl and von Wirén 2014; Gruber et al. 2013). For example, in many plant species, phosphorus (P) starvation results in decreased growth of the main root, increased lateral root formation and stimulation of root hair formation (Alaguero-Cordovilla et al. 2018; Péret et al. 2014). This results in a fundamental reorganization to a more branched and dense root system, while the shoot responds in the opposite way: higher apical dominance (i.e. less branching). Hence, conceptually, phosphate starvation promotes root branching and inhibits shoot branching in an apparent attempt to increase mineral nutrient absorption potential of the root system, and to reduce demand by the shoot.

Here, we take a systematic approach to test the role of SL in root system development with a focus on root branching. We used previously characterized petunia mutants known for their DECREASED APICAL DOMINANCE (dad) phenotype (Drummond et al. 2009; Napoli 1996; Simons et al. 2007; Snowden et al. 2005). These mutants are defective in either SL biosynthesis (*dad1*) or SL sensing (*dad2*) and have previously been shown to encode two highly conserved proteins, the CCD8 enzyme (*DAD1*) (Snowden et al. 2005), and an alpha/beta-hydrolase functioning as an SL receptor (*DAD2*) (Hamiaux et al. 2012), respectively. We show that *dad2* exhibits altered root system architecture, and that this is due mainly to regulating the outgrowth of lateral roots from the main root. The fact that this phenotype is not observed in *dad1* mutants suggests that DAD2 may promote lateral root outgrowth by perceiving a DAD1-independent SL analog (SL-like, further referred to as SLL). This scenario is reminiscent of the SL-related Karrikin (KAR) pathway (Waters and Nelson 2023). In this case, smoke-related signals are perceived by a receptor homologous to DAD2/D14 (Karrikin-insensitive2, KAI2; D14-like). However, the KAR pathway is conserved even in species that are not adapted to cycles of wildfires and recolonization of burned habitats, indicating that in these species, KAI2 may serve as a receptor for yet unknown endogenous hormonal signals (Waters and Nelson 2023).

## Materials and methods

### Plant material and growth conditions

*Petunia hybrida* line V26 was used as the wild type, from which the mutants *dad1* and *dad2* were derived (Napoli and Ruehle 1996). Promoter-Gus lines (*pDAD2::GUS*) where generated with *P. hybrida* W115 Mitchell diploid. In general, seedlings were germinated and grown in 8g/L plant agar (Duchefa; www.duchefa-biochemie.com), supplemented with in ½ strength MS (Duchefa) (Murashige and Skoog 1962). Additional supplement of 1µM of NAA (Sigma) and distilled water was used as a mock for the study of Auxin treatments for a period of 10 days. For phosphate treatments, ½ MS medium was prepared without phosphate as follows: Final concentration of macronutrients (from a 100x stock solution) was 825 mg/l NH_4_NO_3_, 220 mg/l CaCl_2_ · 2H_2_O, 90.35 mg/l MgSO_4_ · 7H_2_O, and 950 mg/l KNO_3_. Micronutrients (applied from a 1000x stock solution) were 3.1 mg/l H_3_BO_3_), 0.013 mg/l CoCl_2_ · 6H_2_O, 13.9 mg/l FeSO_4_ · 7H_2_O), 11.15 mg/l MnSO_4_ · 4H_2_O, 0.42 mg/l KI, 0.13 mg/l Na_2_MoO_4_ · 2H_2_O, 4.3 mg/l ZnSO_4_·7H_2_O, 18.4 mg/l FeNaEDTA, and 0.013 mg/l CuSO_4_ · 5H_2_O. KH_2_PO_4_ was added to this basal medium to reach the indicated concentrations. In general, seedlings were first germinated in ½ MS media for 10 days and only the seedlings with a comparable primary root length were selected for the treatments. All seedlings were vertically grown in 12×12 cm square plates in a growth chamber at 24 °C with a 16 h photoperiod of 32 μmol m^-2^ s^-1^ illumination.

### Cloning

Approximately 2 kb of the *DAD2* promoter region was amplified by PCR with a set of primers (5ʹ-AATGTGGGTGGTCTCAGCTT-3ʹ and 5ʹ-AAATACCACACACCCCTCCA-3ʹ) using genomic DNA from *Petunia hybrida* wild type W115 as a template. The amplified products were cloned into pENTR207 (Invitrogen). The entry vector was mixed with pKGWFS7, and incubated with LR clonase (Invitrogen) to produce *pDAD2::GUS* by an LR reaction. The resulting binary vector was transformed into *Agrobacterium tumefaciens* GV3101 through electroporation for leaf-disc transformation of *Petunia hybrida* wild type W115.

### Leaf disc transformation

Pre-cultures of *A. tumefaciens* GV3101 were prepared from 100 μL frozen glycerol stocks in 5 mL of 10g/L LB Broth (Lennox) (www.huberlab.ch) and adjusted to pH 7.5. Cultures were supplemented with 50 mg/l spectinomycin (Roth; www.carlroth.com) and 25 mg/l gentamycin (Roth) and cultured overnight at 28°C with shaking (250 rpm). Bacterial suspensions with an optical density at 600 nm of 0.6 (OD_600_ = 0.6) were pelleted at 2500 g for 15 min at 4 °C. The supernatant was discarded, and the pellet was re-suspended in 50 mL of half strength liquid MS medium supplemented with 20 μM acetosyringone (Sigma; www.sigmaaldrich.com). Young Petunia leaves were harvested from growth chamber-grown plants, surface-sterilised by soaking in 1.4 % (w/v) sodium hypochlorite for 10 min, and subsequently rinsed five times in sterile water. Young leaf explants were cut to obtain approximately 10 x 10 mm rectangular explants (50-60 pieces) and submerged in the bacterial solution for 20 min.The petunia explants were placed on co-cultivation media for 3 nights in the dark: 2.2 g/l MS medium (Murashige and Skoog, 1962) supplemented with 20 g/l sucrose, 10 g/l glucose, and 8 g/l plant agar (Duchefa). The pH was adjusted to 5.7 before autoclaving and 20 μM acetosyringone was added after autoclaving. The explants were transfered from co-cultivation media to selecting media in 9 cm diameter Petri dishes containing callus- and shoot-inducing medium (pH adjusted to 5.7 before autoclaving), that is a solid regeneration medium based on half strength MS medium (2.2 g/l) supplemented with 20 g/l sucrose, 10 g/l glucose, 8 g/l plant agar (Duchefa). After autoclaving and before pouring, the medium was supplemented with MS vitamins (Sigma), 250 mg/l cefotaxime (Duchefa), 1 mg/l 6-benzylaminopurin (BAP) and 300 mg/l kanamycin (Roth). The explants were transferred into fresh media every 3 weeks. Once shoots had regenerated and reached 2 cm, they were cut from the explant and transferred to root-inducing medium containing half MS medium plus MS vitamins, 250 mg/l cefotaxime and 0.2 mg/l 1-naphthaleneacetic acid (NAA) for 2 days. Next, the NAA-treated shoots were transferred to hormone-free medium for root elongation containing half MS medium plus MS vitamins and 250 mg/l cefotaxime. The shoots were transferred into fresh medium every 3 weeks until they reached a length of 4 to 8 cm and produced a well-developed root system. Then, the plants were acclimatized and transferred to growth chamber conditions. Shoot regeneration and root induction were performed in a growth chamber at 24 °C with a 16 h photoperiod of 32 μmol ^m-2 s-1^ illumination.

### Staining, sectioning, microscopy, and imaging

β-Glucuronidase (*GUS*) staining was performed as described (Sessions et al. 1999) with minor modifications. The samples were submerged in the staining solution (50 mM sodium phosphate buffer pH 7.0, 0.2% Triton-X-100, 10 mM potassium ferrocyanide, 10 mM potassium ferricyanide, 1 mM X-gluc) and vacuum inflitrated for 10 min followed by incubation at 37° C for 12 h. For sectioning, GUS-stained samples were embedded in paraffin in 20/20 mm Tissue-Tek® Cryomold® following established protocols (Zimmermann and Werr 2005) with minor modifications. Paraffin-embedded samples were sectioned using a Reichert-Jung RM2065 microtome with a thickness of 8 µm, processed as described (Feddermann et al. 2010), and mounted in Merckoglas (https://www.sigmaaldrich) for microscopic analysis with a Leica DMR. For cell size measurements, *dad2* and wild type (V26) seedlings were grown for two weeks in ½ strength MS and stained in 0.1% calcofluor white (Cyanamid) for 30 min before analysis on an SP5 confocal microscope. The length of five cells adjacent to LRPs per biological replicate was measured.

### Leaf vein analysis

Seedlings were grown on plant agar (Duchefa), supplemented with ½ strength MS medium. After 2 weeks, leaves were collected, decolorized using 50 % ethanol, and stained with 1% basic fuchsin (Sigma) for 45 mins. After three washes with water the samples were analyzed using a Leica DMR microscope. Images of leaves of similar size were acquired. Then an equivalent field of analysis was defined for each image, and the vein junctions and vein ends were visually counted.

### Microarray analysis

Microarray design, production, hybridization, analysis, and data normalization was performed by NimbleGen as described (Breuillin et al. 2010). Briefly, a 4-plex microarray with 72 000 features was designed using the ArrayScribe software (www.nimblegen.com) to generate 3 optimized independent probes per gene, with an average length of 36 base pairs per probe. Shorter sequences were represented by 2 probes. Comparison and analysis of expression data sets was carried out with Fire2.2 (Garcion and Metraux 2006).

### Statistical Analysis

Statistical analysis was performed in GraphPad Prism 10 as indicated in the legends.

## Results

### General root development and plasticity towards phosphate supply in petunia

Elaboration of the root system involves species-specific developmental programs but is also highly plastic towards environmental conditions. In particular, nutrient status impinges on root development by affecting growth of the primary root, and root branching by lateral root formation (Gruber et al. 2013; López-Bucio et al. 2003; López-Bucio et al. 2002; Pérez-Torres et al. 2008; Williamson et al. 2001). Consequently, root system architecture is influenced by various endogenous and exogenous inputs, and integrates hormonal, nutritional and additional environmental factors. A well-known example of root system plasticity is the response to nutrient starvation, which tends to favors root branching at the expense of primary root elongation, conceivably increasing nutrient uptake capacity by increasing relative surface of the root system (Giehl and von Wirén 2014). This effect has been thoroughly studied in the case of phosphate (P_i_) starvation (López-Bucio et al. 2002).

In order to characterize root system architecture of petunia, and to investigate its response to P_i_ starvation, we grew petunia seedlings in a range of P_i_ concentrations from 20 μM to 2 mM on half-strength MS medium. At concentrations up to 200 μM of KH_2_PO_4_, primary root (PR) elongation during 5 days was minimal (**Fig. 1a,b**), while 660 μM and 2 mM caused a marked increase in PR elongation (**Fig. 1a,b**). This suggests that up to 200 μM of P_i_, plantlets are P-limited. Interestingly, lateral root (LR) density was not significantly affected by P_i_, although the highest P_i_-concentration tended to result in the lowest LR density (**Fig. 1c**). These results indicate that P_i_-limitation severely inhibits PR growth while LR formation is not P_i_-responsive over the tested range of P_i_ concentrations.

**Figure 1.**
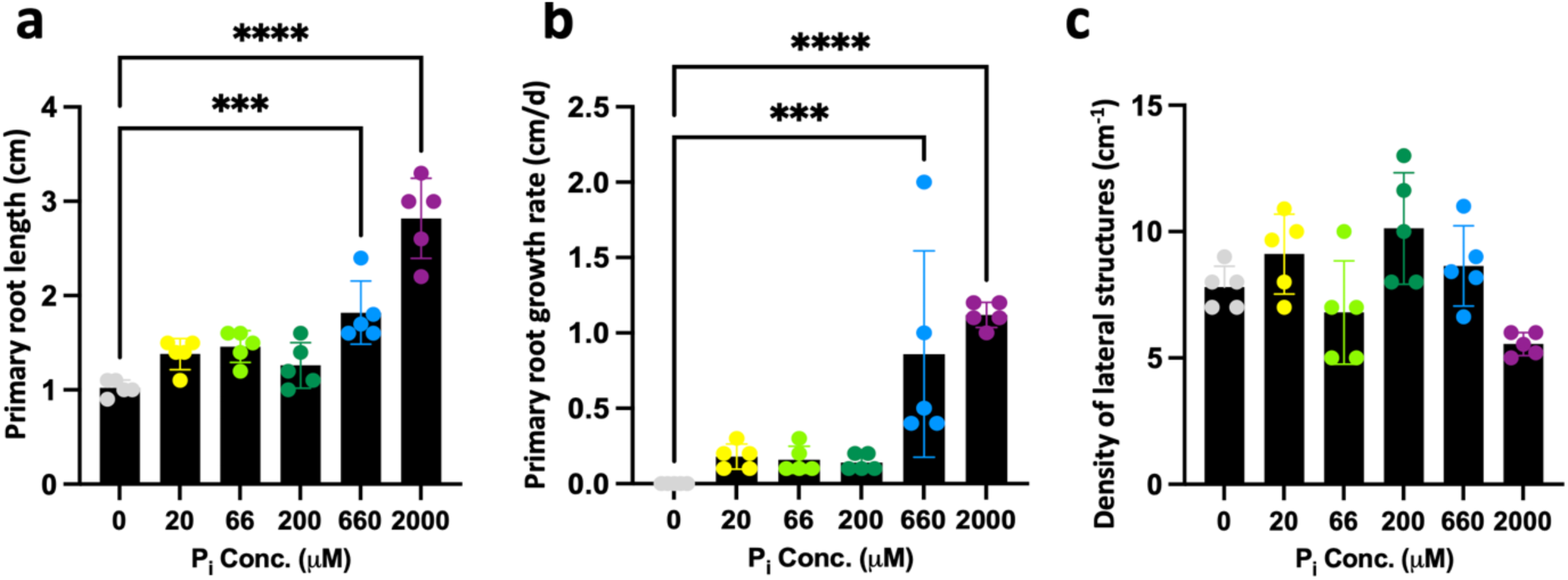
General root development in petunia in response to phosphate status. Primary root length **(a)**, growth rate of primary root **(b)**, and density of root branching sites (lateral roots and lateral root primordia; **(c)**) are shown for various concentrations of KH2PO4 as indicated up to 2 mM. Columns represent the mean (n=5) ± SD (one-way ANOVA and Tukey’s post-hoc test). Significant differences relative to the controls are indicated with asterisks (*** p<0.001; **** p<0.0001).

### *Dad2* grows more slowly than *dad1* or wild type, and meristems are slightly shorter

Changes in plant architecture associated with P_i_-starvation are likely to involve SL (Kapulnik et al. 2011; Koltai 2011; Ruyter-Spira et al. 2013; Ruyter-Spira et al. 2011). For example, reduced shoot branching caused by P_i_-starvation is associated with increased levels of SL biosynthesis, which acts as an inhibitor of shoot branching (Al-Babili and Bouwmeester 2015; Domagalska and Leyser 2011). On the other hand, SL secretion from the roots of P_i_-starved plants promotes colonization by symbiotic arbuscular mycorrhizal (AM) fungi (Kretzschmar et al. 2012), which in turn increase plant nutrition with P_i_ (Smith and Read 2008). SL also impinges on root development, although its role in root system architecture is less clear than in the shoot (Al-Babili and Bouwmeester 2015; Domagalska and Leyser 2011; Ruyter-Spira et al. 2013; Ruyter-Spira et al. 2011).

In order to address the role of SL in root architecture in petunia, we employed SL-related mutants known as *decreased apical dominance1* (*dad1*), and *dad2*, which have been characterized at considerable phenotypic, genetic, and molecular detail (Drummond et al. 2009; Drummond et al. 2012; Hamiaux et al. 2012; Lee et al. 2020; Napoli 1996; Simons et al. 2007; Simons and Ikonen 1997; Snowden and Napoli 2003; Snowden et al. 2005). It is worth noting that these mutants were identified based on excessive shoot branching and subsequent phenotypic analysis was essentially restricted to the shoot phenotype (Drummond et al. 2009; Napoli and Ruehle 1996; Simons et al. 2007; Snowden et al. 2005), hence the role of SL in root development in petunia is unknown.

We assessed root development in *dad1*, which is defective in SL biosynthesis due to a mutation in *carotene cleavage dioxygenase 8* (*CCD8*) (Snowden et al. 2005), and in *dad2*, which has a mutation in an alpha/beta-hydrolase (also known as D14) that acts as an SL receptor (Hamiaux et al. 2012). DAD2 binds SL and interacts with an SCF complex to mediate SL response, in part by mediating degradation of repressors of SL response (known as D53 in rice and SMAX in Arabidopsis) (Lee et al. 2020). The molecular functions of DAD1 (CCD8) and DAD2 (alpha/beta-hydrolase) are well conserved among land plants, including monocots and dicots (Ruyter-Spira et al. 2013; Waters et al. 2017).

Plants were grown in vertical plates on ½ MS medium and general root development was assessed. First, we measured general growth of the primary root (PR). *Dad1* mutants grew with a similar rate and to a similar final length as the wild type (V26), but *dad2* grew significantly slower (**Fig. 2a,b**). Root growth is determined by cell division and cell elongation, hence, we measured the size of the root apical meristem after staining with NBT (Chen et al. 2021). Meristems were generally shorter in *dad2*, while meristem width was not affected (**Fig. 2c-f**). Hence, *dad2* root growth may be slower because the meristem is shorter, allowing for fewer cell divisions to occur in the longitudinal direction.

**Figure 2.**
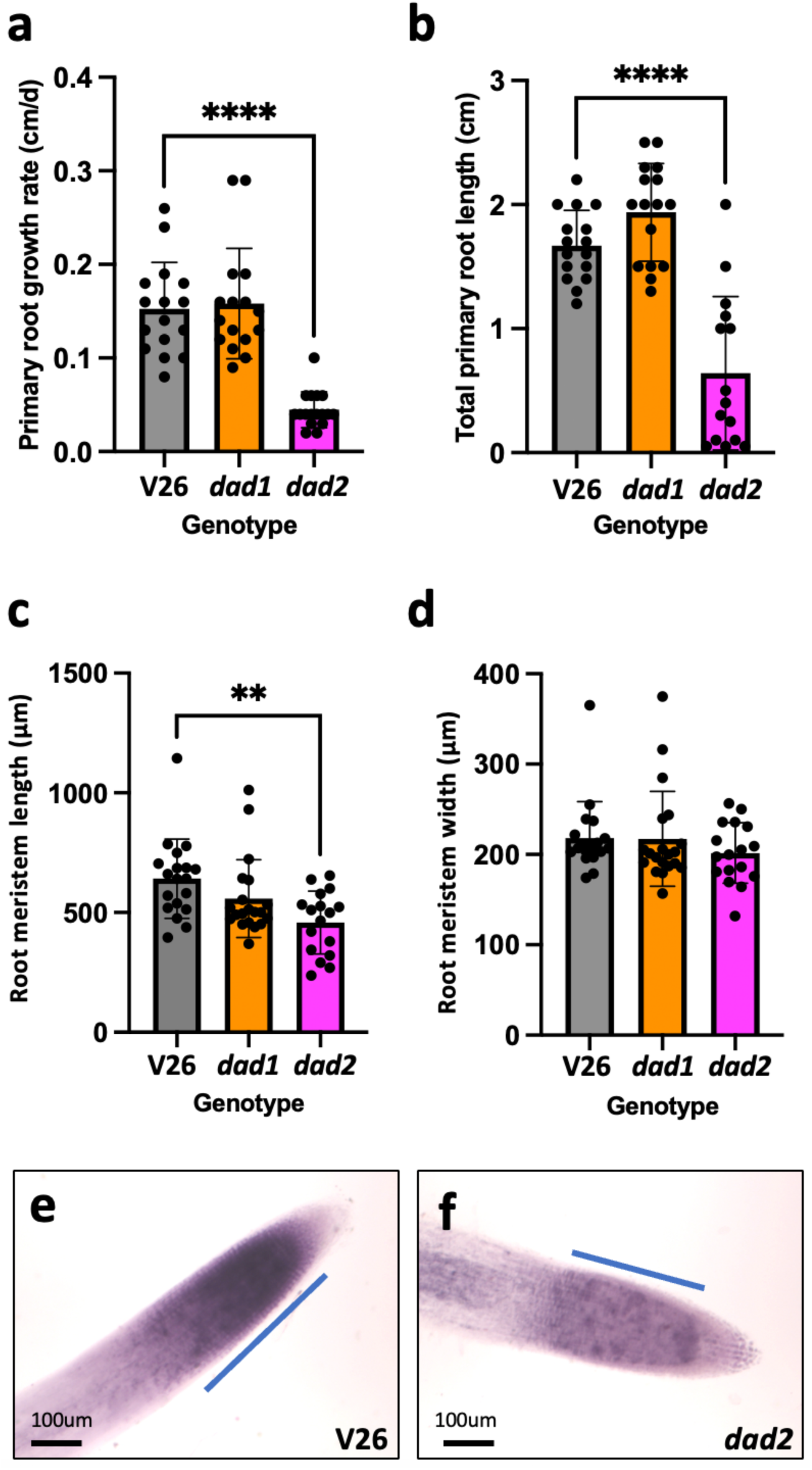
Primary root growth in wild type (V26) and SL-related mutants *dad1* and *dad2*. Primary root growth rate **(a)**, final root length after 1 week **(b)**, root meristem length **(c)**, and root meristem width **(d)** of wild type V26 and the mutants *dad1* and *dad2*, as indicated. **(e,f)** Root meristem dimensions (visualized by NBT staining) of a wild type **(e)**, and a *dad2* mutant **(f)**. Columns represent the mean (n=16) ± SD (one-way ANOVA and Tukey’s post-hoc test). Significant differences relative to the wild-type controls (V26) are indicated with asterisks (** p<0.01; **** p<0.0001). Size bars 100 µm.

### Lateral root primordia are retarded in *dad2*

Next, we explored root system architecture by quantifying root branching. Lateral roots (LR) are formed at a distance from the root tip by cells of the pericycle that undergo dedifferentiation and establishment of new root meristems within the main root. From their point of initiation, lateral roots have to penetrate successively the endodermis, the cortex and the epidermis, before they reach the root surface and emerge into the surrounding soil (Banda et al. 2019). In order to quantify LR initiation, we used NBT staining which highlights meristems at a very early stage, and therefore allows to detect them just after their inception (**Fig. 3a,b**). The total number of initiated LR (including LR primordia and emerged LRs) was similar in all genotypes (**Fig. 3c**). However, *dad2* mutants had more LR primordia (LRPs) (**Fig. 3d**), apparently because a higher percentage of initiated LRs remained within the main root (**Fig. 3e**). This suggests that LR outgrowth requires DAD2 activity.

**Figure 3.**
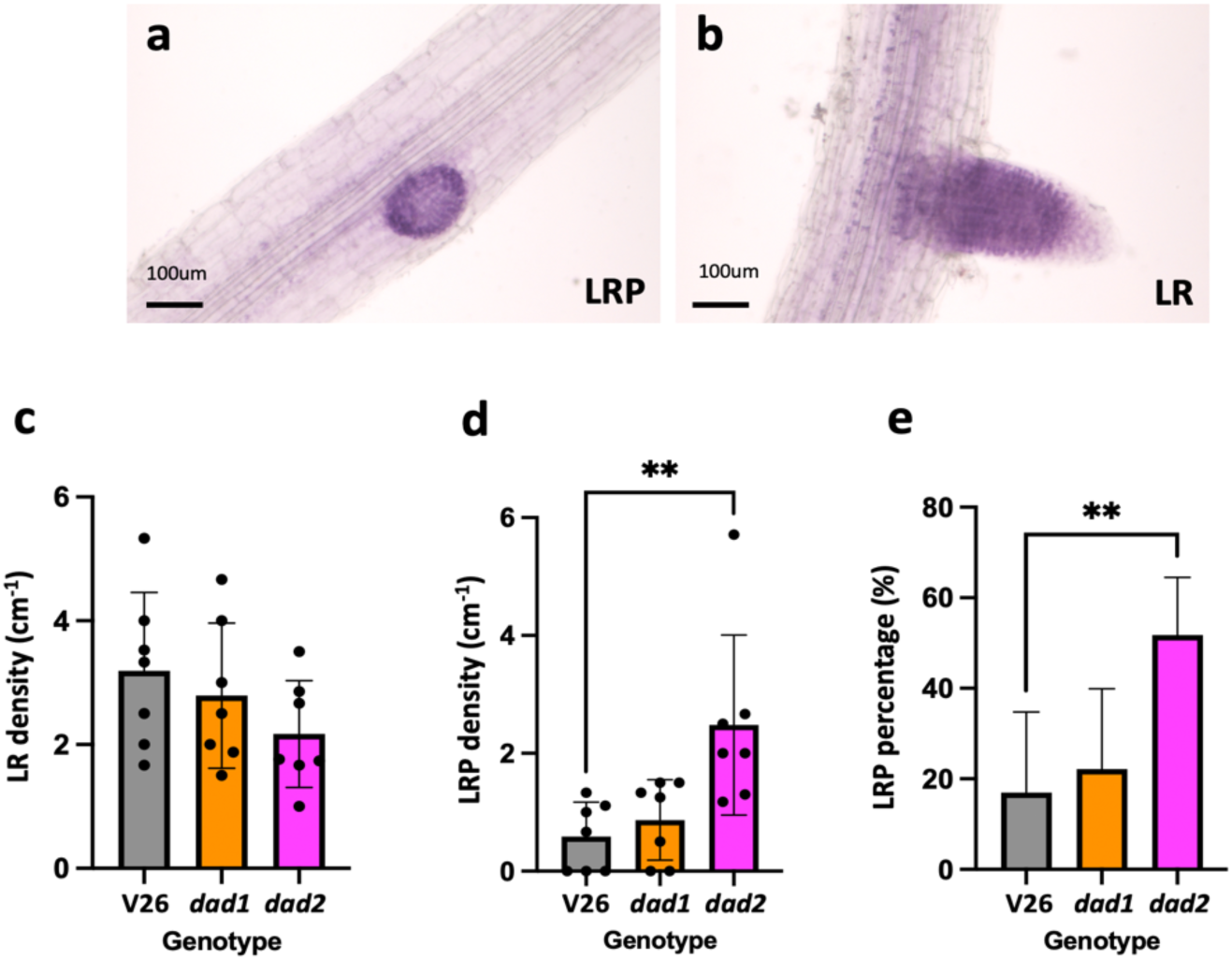
Lateral root formation in *dad1* and *dad2* mutants. Developing branching sites of the main root were classified as lateral root primordia (LRP; **(a)**) or lateral roots (LR; **(b)**), depending on whether they had penetrated the epidermis of the main root. **(c-e)** Density of lateral roots **(c)**; lateral root primordia **(d)**, and percentage of LRPs relative to all lateral structures **(e)** are shown. Columns represent the mean (n=7) ± SD (one-way ANOVA and Tukey’s post-hoc test). Significant differences relative to the wild-type controls (V26) are indicated with asterisks (** p<0.01). Size bars 100 µm.

### Cell growth around LR primordia is not affected in dad2 mutants

LRP emergence in *dad2* mutants could potentially be retarded due to defects in cellular growth and/or organization, hence, we measured cell length around LRPs from confocal pictures of calcofluor-stained roots (**Fig. S1a,b**). Although *dad2* cells tended to be slightly shorter, the difference was not significant (**Fig. S1c**), indicating that cell growth and organization is not affected in the mutant roots.

### The *DAD2* promoter is active along the vasculature and at sites of lateral root emergence

Receptors often mediate developmental responses in a cell-autonomous fashion, since they usually mediate intracellular signaling events that have direct consequences for the perceiving cell (with the additional possibility of secondary effects on other cells). Hence, we suspected that *DAD2* might be expressed in roots to mediate the developmental effects observed in the *dad2* mutant. In order to reveal the expression pattern of *DAD2* with cellular resolution, petunia was transformed with a *pDAD2::GUS* construct.

Transgenic *pDAD::GUS* plants expressed the GUS marker immediately after germination (**Fig. 4a**), in particular in the shoot apex and the cotyledons, where expression was patchy, but often associated with the vasculature. At later stages of development, *pDAD2::GUS* signal became weaker, with expression becoming progressively confined to the vasculature and the hydathodes at the tips of the cotyledons, and in the young leaves (**Fig. 4b**). Subsequently, GUS signal was observed throughout the shoot, most prominently in the apical parts of the stem and along the vasculature (**Fig. S2a,b**). In leaves, local signal was occasionally elevated in the vasculature at branches and ends of vascular strands (**Fig. 4c,d**). In sectioned stem tissues, the signal was most prominent inside and outside the wood in cells that may represent xylem and/or phloem parenchyma (**Fig. S2c,d**).

**Figure 4.**
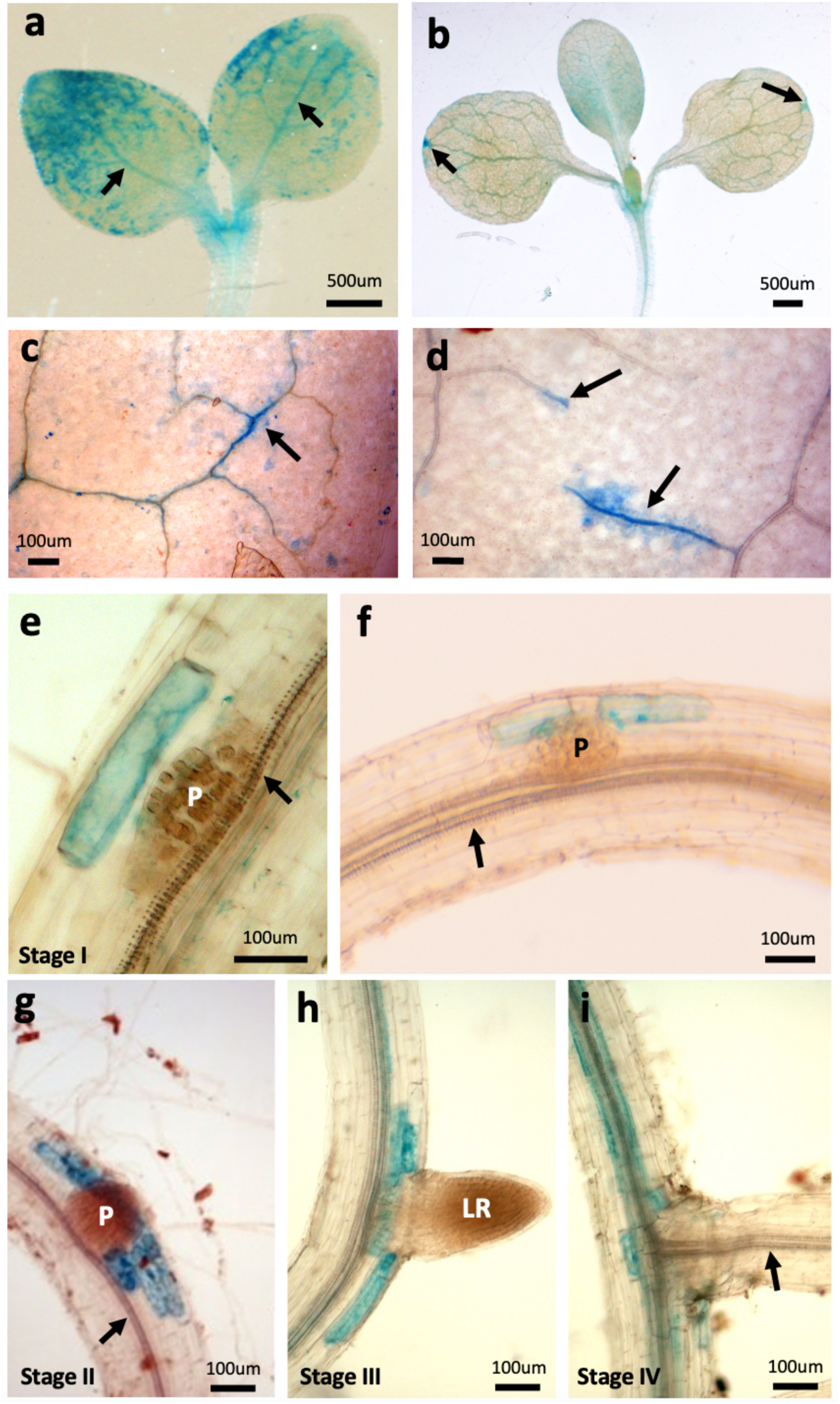
Expression pattern of pDAD2::GUS during seedling development. Seedlings transformed with the *pDAD2::GUS* construct were stained for glucuronidase activity and destained for analysis. (a) young seedling just after germination with elevated staining along vascular strands (arrows), (b) seedling with fully expanded cotyledons and the first true leaf. Note elevated staining at cotyledon leaf tips (arrows). (c-d) staining in the vasculature of fully expanded true leaves.

In young roots, *pDAD2::GUS* was hardly detectable, however, at sites of LR formation, the *DAD2* promoter became markedly induced in just one or two cells adjacent to the young LRPs (**Fig. 4e,f**). At later stages of LRP development, the GUS expression domain became extended towards inner and outer cell layers, but GUS-expressing cells were always in direct contact to the developing LR primordium during emergence of the LR through the epidermal layer (**Fig. 4g,h**). When the lateral root was fully functional, as evidenced by the continuous vascular system between main root and lateral root, expression at the base of the lateral roots became weaker (**Fig. 4i**). At this time, general expression along the vasculature of the main root became induced, conceivably reflecting general root maturation (**Fig. 4i, Fig. S2e**). In cases where lateral roots were induced in more mature regions of the root, the same pattern of *pDAD2::GUS* induction was observed in individual cortex cells above LRPs (**Fig. S3**). Sections from paraffin-embedded mature roots revealed signal surrounding the central woody stele (**Fig. S4a,b**), and cells interspersed in the wood that may represent xylem parenchyma (**Fig. S4c,d**).

### Vasculature is denser in leaves of *dad2* mutants than in the wild type or *dad1*

Since *pDAD2::GUS* expression was elevated along the leaf vasculature (**Fig. 4c,d**), we tested whether *DAD2* has a role in vascular pattering. Leaves of *dad2* mutants were stained with basic fuchsin which associates with lignin and therefore highlights xylem strands (Kapp et al. 2015). As distinct features of leaf vascular patterns, we counted vascular branches and vein ends of the leaf vasculature system. We found that the density of both, branches and vein ends, were increased in the SL sensing mutant *dad2*, but not in the SL biosynthetic mutant *dad1* (**Fig. 5a,b**). These results indicate that DAD2 perceives an SL-related (but CCD8-independent) signal that attenuates the formation of vascular strands in the leaves.

**Figure 5.**
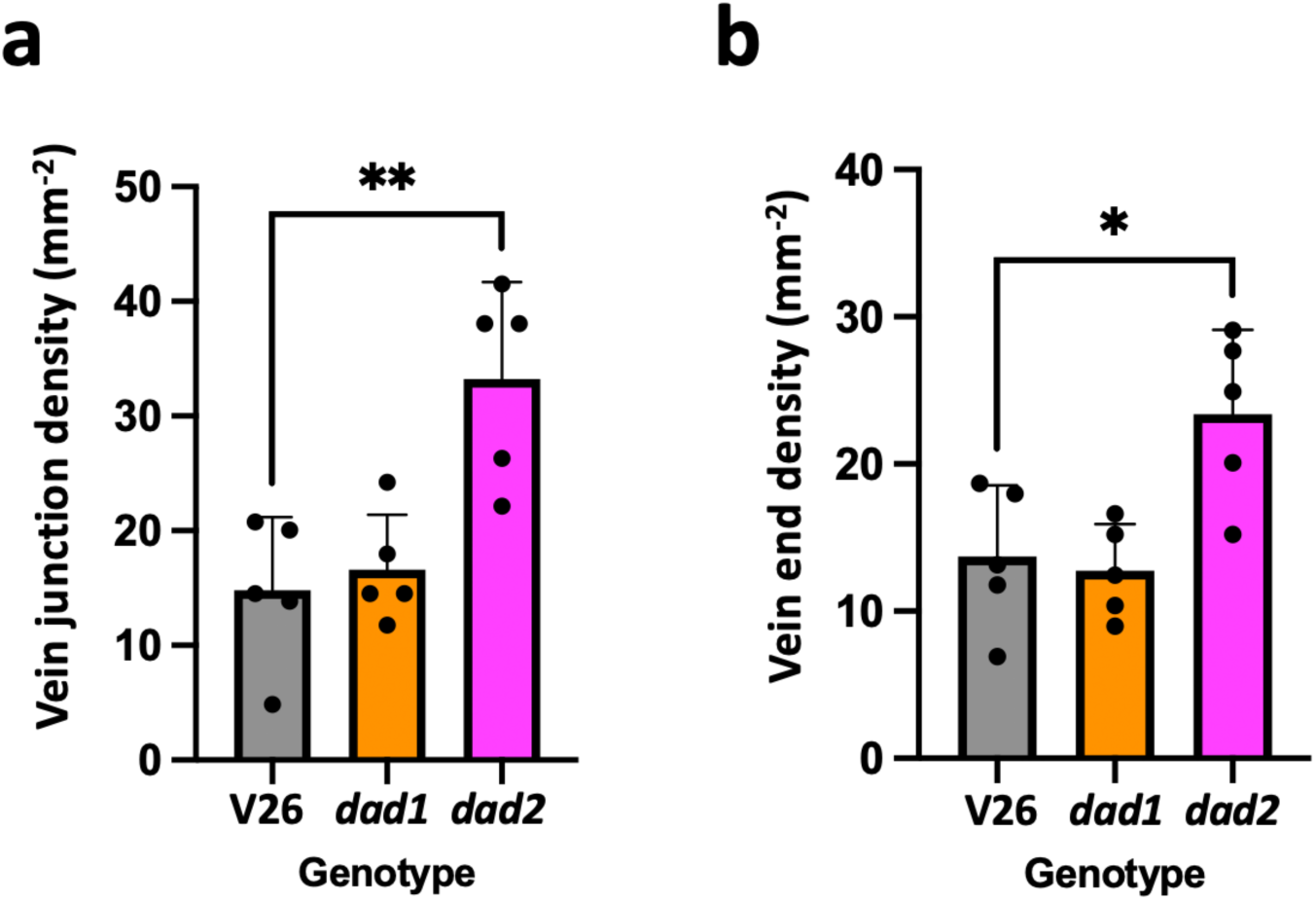
Leaf venation phenotype of *dad1* and *dad2* mutants. Fully expanded leaves of wild type V26, *dad1*, and *dad2* mutants were stained with basic fuchsin to reveal vasculature and visually assessed to determine the number of vein junctions **(a)** and vein ends **(b)** in equivalent regions in the center of fully expanded leaves. Columns represent the mean (n=5) ± SD (one-way ANOVA and Tukey’s post-hoc test). Significant differences relative to the wild-type controls (V26) are indicated with asterisks (* p<0.05; ** p<0.01).

### The LRP emergence phenotype in *dad2* mutants is complemented by NAA

Root development and LR formation are sensitive to auxin and require polar auxin transport (PAT). Auxin inhibits root elongation and root growth rate, while it stimulates the initiation of LRP (Cavallari et al. 2021; Roychoudhry and Kepinski 2022). PAT is required both, for maintenance of an active root meristem (Sabatini et al. 1999), and for LR initiation (Benkova et al. 2003). SL controls auxin flux in the plant by regulating the subcellular localization of auxin transporters of the PIN-FORMED (PIN) family (Zhang et al. 2020; Shinohara et al. 2013; Domagalska and Leyser 2011; Hayward et al. 2009), thereby providing an integrated auxin transport network that coordinates meristem activity across the entire shoot (van Rongen et al. 2019). On the other hand, SL acts downstream of auxin, since its biosynthesis is stimulated by auxin (Mashiguchi et al. 2021). Hence, the interactions between SL and auxin are complex. In order to test whether the root phenotypes in *dad2* mutants are related to auxin, we tested the response of *dad1* and *dad2* mutants towards the synthetic auxin naphthylacetic acid (NAA), applied at a concentration of 1 µM, which was the minimal concentration required for consistent LR induction (**Fig. S5**). As expected, NAA inhibited PR growth in all genotypes (data not shown), while it stimulated induction of LRP in all genotypes (**Fig. 6a**). Interestingly, the fraction of root primordia in *dad2* was reduced by NAA to reach similar levels as in the other genotypes, indicating that exogenous auxin can restore the LRP emergence phenotype of *dad2* by promoting primordium outgrowth (**Fig. 6b**). These results indicate that auxin mediates LRP outgrowth downstream of *DAD2* and that outgrowth as such is not affected in *dad2* mutants if sufficient auxin is supplied to the roots.

**Figure 6.**
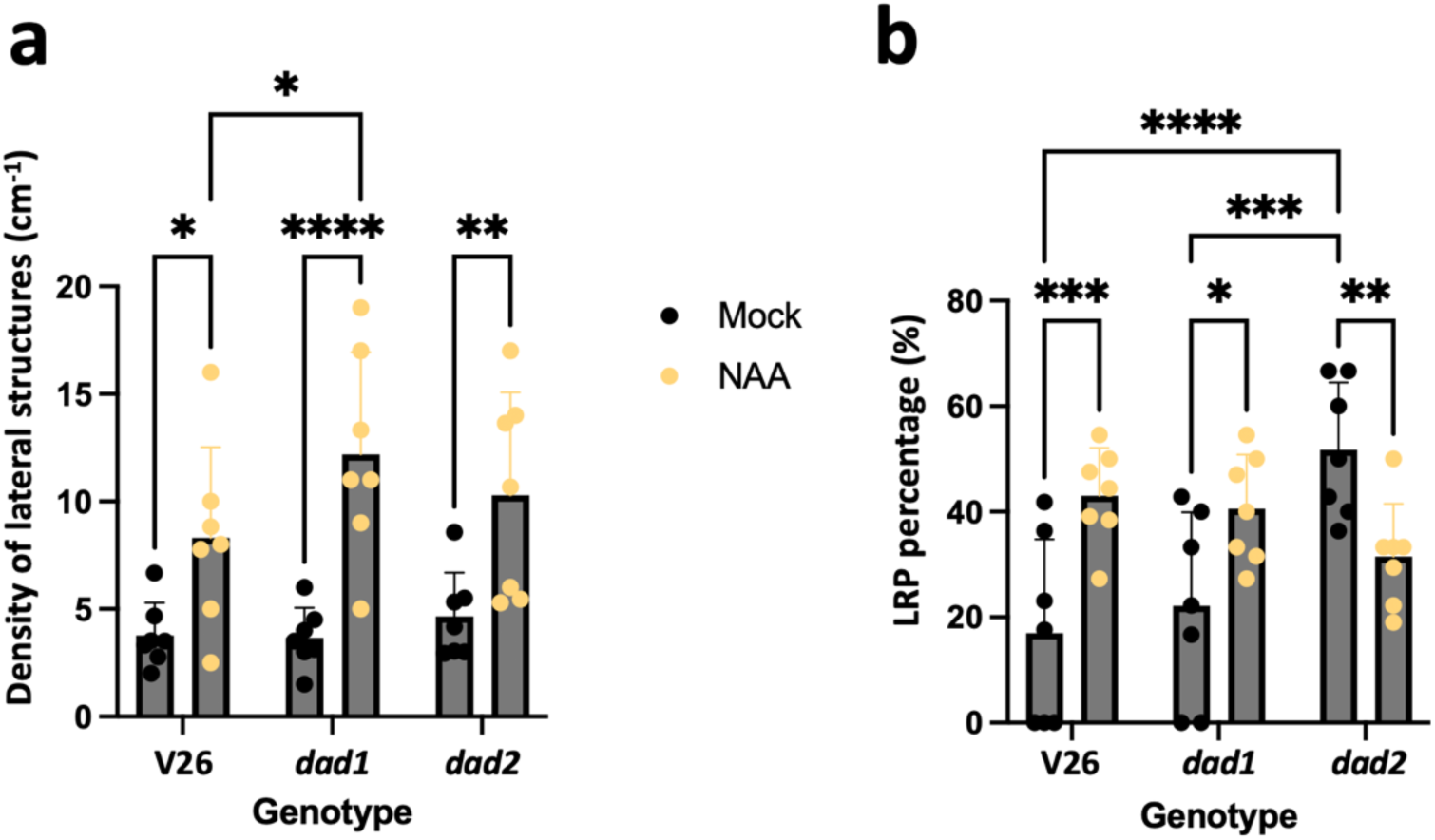
Recovery of the root emergence phenotype in *dad2* by auxin supply. Plantlets of wild type V26, *dad1*, and *dad2* mutants were grown on ½ MS plates with 1 µM NAA for 5 days. The density of root branching sites (lateral roots and lateral root primordia) **(a)** and the percentage of lateral root primordia among all branching sites **(b)** is shown. Columns represent the mean (n=7) ± SD (one-way ANOVA and Tukey’s post-hoc test). Significant differences relative to the wild-type controls (V26), or relative to untreated controls (Mock) are indicated with asterisks (* p<0.05; ** p<0.01; *** p<0.001; **** p<0.0001).

### SL- and auxin-inducible genes in petunia roots

Since DAD2 function is required for LRP outgrowth (**Fig. 3**), a phenomenon that is also under the control of auxin and downstream cell wall remodeling proteins (Swarup et al. 2008), we asked what genes are induced by SL in the root system, and whether SL-inducible genes are also responsive to auxin. We used previously described microarrays (Breuillin et al. 2010) to explore the transcriptomic response of soil-grown plants towards the synthetic SL analogue GR24, and to assess the effect of auxin and gibberellin (GA3; regarded here as an unrelated growth hormone for comparison) on the expression of SL-responsive genes in petunia roots. Among 24’618 gene IDs represented in triplicates on a Nimblegen microarray (Breuillin et al. 2010), nine gene IDs were induced by GR24 at least 3-fold in two independent experiments (**Table S1**). They can be assigned to four main functional groups of genes: i.) three glutathione S-transferases (GSTs), ii.) three cytoskeleton- and cell wall-related genes (tubulin, an arabinogalactan protein, and the lignin biosynthetic enzyme caffeoyl-CoA-3-O-methyltransferase), iii.) a phosphate signaling component (SPX protein), and, notably, iv.) an auxin transport protein (PILs7). It belongs to a PIN-related clade (PIN-likes; PILS) that controls intracellular auxin transport and is involved in intracellular auxin accumulation (Sauer and Kleine-Vehn 2019; Beziat et al. 2017; Feraru et al. 2012; Barbez et al. 2012). Strikingly, four of the mentioned genes were also strongly induced by auxin (the three GSTs, and PILS7), while all nine genes were only weakly affected by the growth hormone GA, used here as an unrelated hormonal control treatment (**Table S1**). These results suggest that an SL-inducible program could act synergistically with auxin transport and auxin-related gene expression to regulate root architecture.

### *pDAD2::GUS* is expressed in a subset of DR5-responsive cells

The expression pattern of pDAD2::GUS along the vasculature (**Fig. 4i, Fig. S2-S4**) coincides with several SL-related genes in Arabidopsis (Khuvung et al. 2022), and it is somewhat reminiscent of auxin markers such as *DR5::GUS* (Mattsson et al. 2003), although *pDR5::GUS* expression decreases with maturation of vascular strands (Mattsson et al. 2003), whereas expression of *pDAD2::GUS* is low increases in mature vascular strands, at least in the root (**Fig. S2e**). In addition, cells adjacent to LRP are known to have increased auxin levels as part of a program related to cell wall loosening during LR emergence (Swarup et al. 2008). To investigate a potential overlap between the *DAD2* expression domain and elevated auxin levels, we crossed *pDAD2::GUS* with a petunia line expressing the auxin marker *DR5::NLS-3xVENUS*, the latter being localized to nuclei due to a nuclear localization signal (Rodriguez-Villalon et al. 2015). Since the analysis of VENUS expression is not compatible with GUS staining, we performed sequential imaging, first for VENUS and then for GUS activity. Live roots were immobilized in low-melting-point agarose and the GFP signal was assessed by confocal microscopy. Then the samples were stained for GUS activity to reveal *pDAD2* expression pattern.

In regions without LRPs, DR5-dependent VENUS signal was weak throughout the root, except for cells along the vasculature with elongated nuclei that exhibited elevated DR5 signal (**Fig. S6a**). Early LRP were characterized by strong DR5 signal (**Fig. S6b**), and in addition, a region around the LRP that corresponds roughly to one cell diameter (ca. 100 µm) was characterized by strong DR5 signal (**Fig. S6b**). At later stages of LR formation, the site of LR emergence continued to show elevated DR5 activity (**Fig. S7**), indicating that cells involved in passage of the LRP experience long-term auxin signaling. Assessing the overlap between the *pDAD2::GUS* and *pDR5::NLS-3xVENUS* expression domains revealed that GUS-expressing cells consistently showed a strong VENUS signal, while only a fraction of VENUS-positive cells coincided with elevated GUS expression (**Fig. 7**). This suggests that DAD2-expressing cells represent a subset of the DR5-positive cells around emerging LRPs.

**Figure 7.**
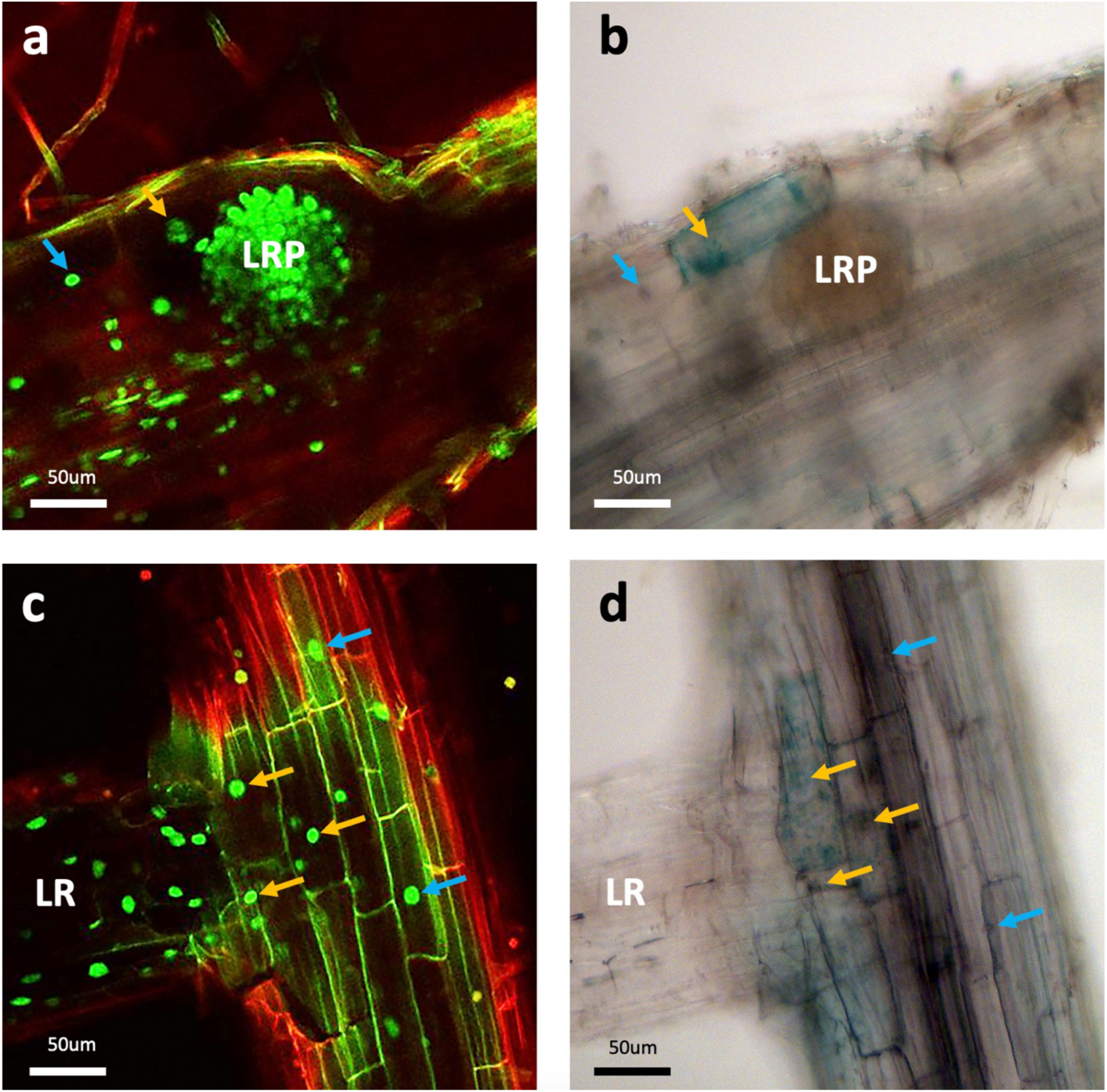
Expression of *pDAD2::GUS* and *DR5::3xVENUS-NLS* at root branching sites. **(a,b)** A lateral root primordium at an intermediate stage of development (compare with Fig. 3a), imaged first alive by confocal laser scanning microscopy **(a)**, and then after GUS staining by bright field microscopy **(b)**. **(c,d)** AN emerged fully grown lateral root imaged first alive by confocal laser scanning microscopy **(c)**, and then after GUS staining by bright field microscopy **(d)**. Arrows indicate DR5-positive cells that express also pDAD2::GUS (orange arrows), or cells that express *DR5::3xVenus-NLS*, but not *pDAD2::GUS* (blue arrows). LR, Lateral root; LRP, Lateral root primordium. Size bars 50 µm.

## Discussion

### Regulation of overall plant growth and P-starvation response by strigolactone

Strigolactone integrates both exogenous (nutritional) and endogenous (developmental) cues to coordinate overall plant growth and architecture. The main function of SL is to promote apical dominance (AD), a developmental program that promotes growth and proliferation of the main shoot, while axillary buds remain inactive (Domagalska and Leyser 2011). Through its responsiveness to P status (Kumar et al. 2015), SL connects P starvation response with plant architecture by inhibiting shoot branching (promoting apical dominance), thereby preventing dilution of scarce P resources over multiple lateral branches. In addition, SL released from the roots promotes the interaction with arbuscular mycorrhizal (AM) fungi (Akiyama et al. 2005; Besserer et al. 2006), which colonize the roots of most plants and supply their host with P_i_ (Smith and Read 2008).

Before SL was recognized as an inhibitor of shoot branching, auxin was held to be the main mediator of AD (Beveridge et al. 2023). In recent years, it became apparent that auxin and SL act in concert to inhibit shoot branching. A well-supported model of apical dominance and shoot branching posits that SL acts mainly (but not exclusively!) by inhibiting polar auxin transport and canalization in axillary buds along the stem (Zhang et al. 2020; Shinohara et al. 2013; Domagalska and Leyser 2011; Crawford et al. 2010; Hayward et al. 2009). Although SL also influences root development (Kapulnik et al. 2011; Koltai 2011; Kumar et al. 2015; Ruyter-Spira et al. 2011), the role of SL in root growth and branching is less well understood, in particular, the root phenotypes of SL-related mutants have been assessed only in a few cases. SL has been reported to stimulate (Koltai 2011; Ma et al. 2020; Ruyter-Spira et al. 2011), or to inhibit (Kapulnik et al. 2011; Koltai 2011; Ruyter-Spira et al. 2011) root branching, depending on the plant species and the growth conditions. Hence, the function of SL in root architecture may be less conserved than its role in shoot branching. Nevertheless, reduced primary root growth appears to be a common feature of SL-related mutants (Arite et al. 2012; Kapulnik et al. 2011; Ruyter-Spira et al. 2011; Sun et al. 2016; Villaécija-Aguilar et al. 2022).

### Petunia root architecture and responsiveness to phosphate starvation

Since root system architecture is influenced by P status (which in turn regulates SL biosynthesis), we first characterized root development over a range of Pi levels up to 2 mM. This experiment showed that limiting P_i_ levels (200 µM or less) resulted in highly reduced primary root growth (**Fig. 1a,b**). Whether this is a mere starvation symptome or a regulated attenuation of primary root growth is difficult to distinguish. On the other hand, LR density was not significantly affected by P_i_ levels (**Fig. 1c**), suggesting that root system remodeling in petunia is less P_i_-sensitive than in *A. thaliana* (López-Bucio et al. 2002; Williamson et al. 2001). In contrast, the effects of exogenous auxin on the root system were as expected: primary root growth was reduced, while LR density was increased by auxin (**Fig. S5a,b**). These results reflect similar findings in various plant species, including monocots (McSteen 2010), and the dicot *A. thaliana* (Roychoudhry and Kepinski 2022), suggesting that the role of auxin in regulation of root architecture is rather conserved.

### A CCD8-independent signal regulates development of petunia through DAD2

In order to explore the role of SL signaling in root system architecture, we chose two well-characterized SL-related mutants in petunia: *dad1* with a defect in SL biosynthesis, and *dad2* which is defective in SL sensing. A striking finding of this study was that *dad2* mutants had specific defects in lateral root emergence and leaf venation, which were not observed in *dad1* mutants (**Fig. 2,3**,**5**). This suggests that DAD2 can perceive DAD1/CCD8-independent signals. In this context, it is interesting to note that SL signaling has a closely related sister pathway known as Karrikin (KAR) signaling pathway (Machin et al. 2020). Karrikin is produced by incomplete burning of plant material and is generated during wildfires (Chiwocha et al. 2009). It serves as a signal in plants adapted to germinate after wildfires, however, also other plants such as Arabidopsis have a KAR signaling pathway, indicating that the KAR receptor KARRIKIN INSENSITIVE2 (KAI2) also perceives endogenous KAI2 ligands (KL), that have so far remained elusive (Waters and Nelson 2023). SL and KAR are both butenolide compounds (Machin et al. 2020), and they both act through similar signaling pathways that share a central component, MAX2 in the ubiquitin ligase complex SCF^MAX2^ (Waters et al. 2017), thus there is potential for interactions between the pathways. Indeed, crosstalk between SL and KAR signaling has been established in Arabidopsis (Li et al. 2022; Machin et al. 2020) and rice (Zheng et al. 2020). In addition, recent research has identified CCD8-independent SL-like signals that are perceived by D14 (S. Al-Babili, personal communication), hence DAD1-independent effects that require DAD2 in petunia could involve such non-canonical DAD2 ligands that mimick SL.

### General expression pattern of pDAD2::GUS

In order to locate presumed sites of SL perception and action, we considered mainly the D14/DAD2 expression pattern in petunia. This is because all biosynthetic genes act on a mobile signal (SL), and grafting experiments have shown that various parts of the plant can be the source of sufficient SL to mediate apical dominance (Kameoka and Kyozuka 2018). Therefore, the expression pattern of SL biosynthetic genes is less relevant for spatial aspects of SL action than the expression pattern of the SL receptor. SL is perceived by a D14/DAD2-based receptor complex, which triggers the degradation of a transcriptional repressor (D53), thereby de-repressing SL-regulated genes (Waters et al. 2017). Based on this scenario, D14/DAD2 is likely to act cell autonomously (although secondary non-cell-autonomous mechanisms cannot be ruled out), hence, SL action is likely to coincide with expression of *D14/DAD2* rather than of SL biosynthetic genes.

### Function of DAD2 and auxin in root emergence

The defect of *dad2* mutants in lateral root emergence (**Fig. 3**), and the specific induction of *DAD2* in cells adjacent to LRP (**Fig. 4e-g**) suggest that DAD2 signaling activates a cell wall-remodeling program required for penetration of LRP through the endodermal, cortical, and epidermal cell layers of the root. Emergence of LRP is regulated by accumulation of auxin which induces cell-wall-remodeling (CWR) enzymes in cells directly above LRPs (Swarup et al. 2008). How could DAD2 activity tie in with this phenomenon?

Inhibition of auxin transport capacity by SL signaling (either through reduced PIN gene expression or by inhibiting PIN protein recycling towards the plasma membrane) would be expected to interfere with auxin accumulation required for LRP inception. This may be the basis of the inhibitory effect of SL on LR formation (Kapulnik et al. 2011; Koltai 2011; Ruyter-Spira et al. 2011). Interestingly, DAD2 is not expressed in young roots before LR initiation, hence SL signaling could not possibly interfere with PIN-dependent auxin accumulation during the initial stages of LR primordium initiation. During LR outgrowth, however, DAD2 becomes induced locally and exclusively in cortical cells adjacent to the LR primordia. It is tempting to speculate that in this context, reduced auxin export due to inhibited PIN activity could reinforce auxin accumulation in these cells caused by auxin importers such as AUX1/LAX3 (Swarup et al. 2008) (see below).

### The challenge to accumulate auxin in a single cell

Since the auxin perception and signaling machinery is localized intracellularly (Wang and Estelle 2014), auxin action requires accumulation within cells. It appears therefore somewhat paradoxical that the auxin transporters (PIN proteins) that regulate most developmental processes involving auxin accumulation are auxin exporters, rather than auxin importers. By principle, they mediate auxin accumulation not in the cells that express these transporters, but in their neighbors. In a developmental context, e.g. in the shoot apical meristem or the root meristem, this requires cell populations, among which auxin is transported against an auxin gradient to accumulate in the center of the population (Sabatini et al. 1999; Reinhardt et al. 2003; Heisler et al. 2005; Jönsson et al. 2006; Smith et al. 2006). Hence, PIN-mediated auxin accumulation is – *per se* – a non-cell-autonomous mechanism that is regulated at a supracellular level.

Given the biochemical function of PIN proteins, auxin accumulation in a single cell can – by definition – not be achieved by PIN induction in this same cell. In contrast, PIN activity in a single cell can only result in auxin depletion in this cell. Instead, SL-mediated cell-autonomous reduction of PIN protein activity (e.g. by reducing PIN protein levels at the plasma membrane) could potentially support auxin accumulation, as it is observed in single cells next to outgrowing LRP (Swarup et al. 2008). SL can reduce the proportion of PIN protein at the plasmalemma (by increased internalization) (Shinohara et al. 2013; Zhang et al. 2020), thereby contributing to cell-autonomous auxin accumulation. Such a role for SL could be important since auxin tends to have the opposite effect by promoting PIN protein accumulation in the plasma membrane (Bennett et al. 2016; Zhang et al. 2020).

### Interaction of SL signaling and auxin at sites of LPR emergence

Taken together, DAD2-mediated signaling could considerably increase auxin accumulation in individual cortex cells that are known to promote LRP outgrowth (Swarup et al. 2008) (**Fig. 8**). An additional contribution could come from induction of PIN-like proteins such as PILS7 (**Table S1**). PILS proteins are auxin transporters that accumulate auxin in intracellular storage compartments such as the ER (Sauer and Kleine-Vehn 2019), hence, PILS7 could potentiate intracellular auxin accumulation in a cell-autonomous manner. This scenario would also be consistent with the finding that in petunia, GR24 activates auxin marker genes such as glutathione-S-transferase (GST) and auxin-responsive cell-wall-remodeling genes (**Table S1**). Finally, an upstream role of SL onto auxin accumulation would also explain the fact that exogenous auxin can rescue LRP emergence in *dad2* mutants (**Fig. 6**). It is interesting to note that in a non-targeted GWAS study in Arabidopsis, PIN-LIKE7 (PILS7) was identified as a determinant that influences shoot-to-root ratio (Deja-Muylle et al. 2022). Strikingly, PILS proteins promote auxin accumulation in cells of Arabidopsis (Barbez et al. 2012), thus providing additional support for the notion that DAD2-expressing cortex cells accumulate auxin (**Fig. 7**).

**Figure 8.**
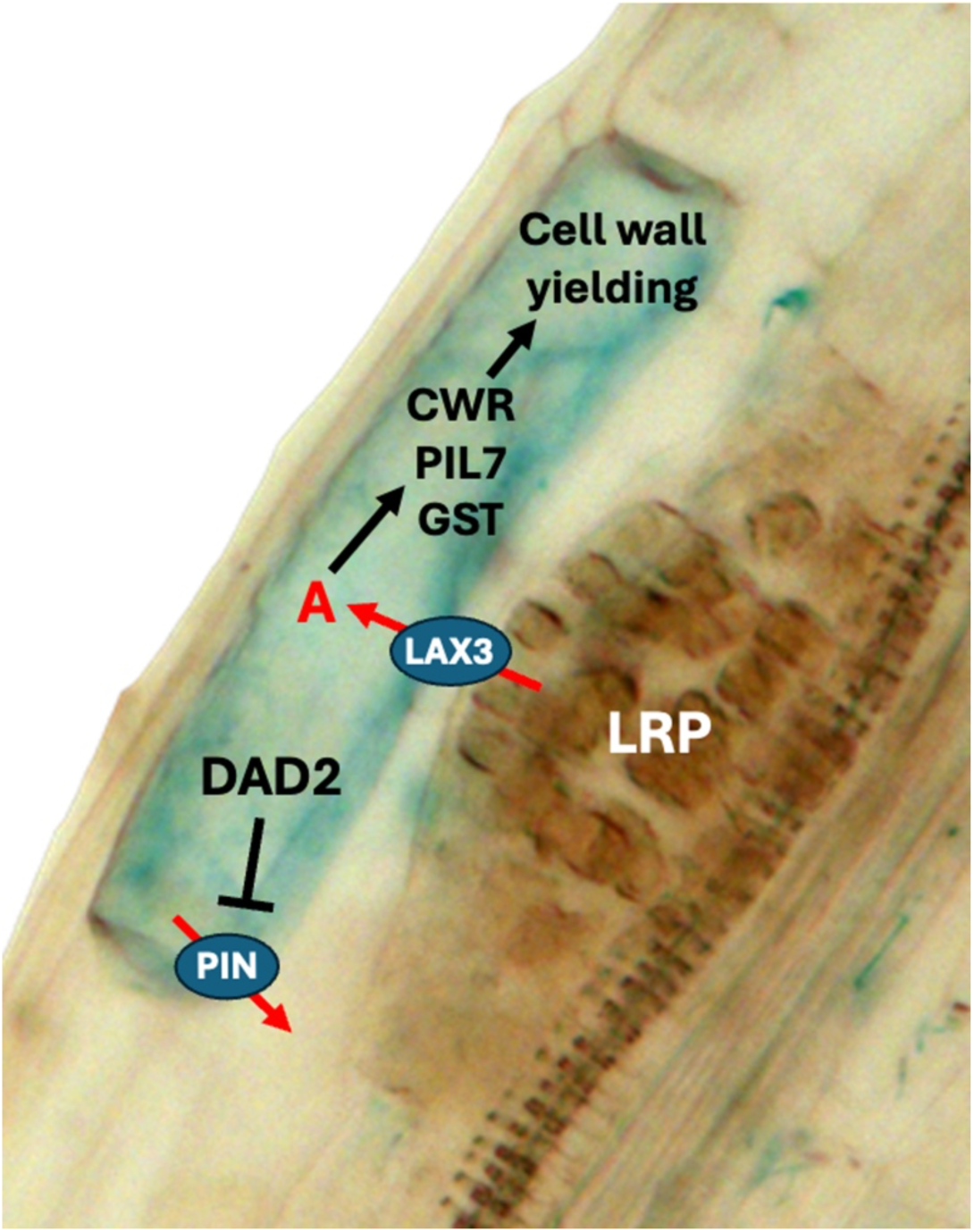
Model of the mode of action of DAD2 and auxin in lateral root formation. Early stage of lateral root formation with a young lateral root primordium (LRP), and an overlaying cortex cell that expresses *DAD2* (blue cell). Auxin imported from the lateral root primordium through LAX3 (red arrow) gets further accumulated by DAD2-dependent inhibition of PIN function. Auxin activates cell wall remodeling enzymes (CWR), and auxin markers such as glutathione-S-transferase (GST) and Pin-Like7 (PIL7). Activation of CWR enzymes results in cell wall yielding and allows the LRP to penetrate through the outer cell layers of the root.

Auxin and SL(L) act in mechanistically different ways, the former by focusing the levels of the active molecule in one/few cells, and the latter by focusing the receptor for SL in the very same cells (**Fig. 7,8**). Through the resulting positive feed-forward mechanism of SL onto auxin accumulation, a subset of individual cells in a population of DR5-responsive cells can become fully committed to cell wall remodeling (to allow LR primordia to pass), without interfering with neighboring cortex cells that should remain unaffected by cell-wall-remodeling agents in order to assure root integrity in the wider context of emerging LRPs.

## Acknowledgements

We thank Francesca Quattrocchio and Pamela Strazzer for providing the *Petunia hybrida* W115 line expressing the *DR5::3xVENUS-NLS* construct, and Kim Snowden for providing seeds of the petunia mutants *dad1* and *dad2*.

## Author contributions

DR conceived the project; KK, PEC, and DR performed experiments and data analysis; KK, PEC, and DR were involved in the writing of the manuscript

## Figure Legends

**Figure S1. Cell size in the vicinity of lateral root primordia in *dad2* mutants.**

**(a)** A lateral root primordium shortly before penetration of the epidermal cell layer imaged by confocal microscopy; **(b)** Same image as in (a) represented as a reverse black-and-white image with two colored epidermal cells. **(c)** Length of cells adjacent to lateral root primordia in wild type (V26) and *dad2* mutants was measured from images as in (b). No significant difference was observed between the genotypes (n=30). Size bars 50 µm.

**Figure S2. Expression of *pDAD2::GUS* during later vegetative growth.**

Seedlings transformed with the *pDAD2::GUS* construct were stained for glucuronidase activity and imaged after removal of chlorophyll. **(a)** Seedling with two cotyledons (cot) and two fully expanded true leaves (only petioles visible, **(b)** seedling with two cotyledons and four fully expanded true leaves. Note weak general expression throughout the shoot and elevated expression along vascular strands. **(c,d)** cross-section (c) and longitudinal section (d) of a stem segment with secondary wood formation (asterisks). Sections in (c,d) were counterstained with phloroglucinol to visualize lignified cells (red). In the stem, GUS signal was observed mainly along vascular strands (arrows in (c,d)) inside and outside of the wood (c), and between files of lignified cells (d). **(e)** In the root system, general GUS signal was observed mainly along the stele of the primary root (arrow) at a distance from the root meristem, while lateral roots exhibited only weak staining in their older proximal parts. Size bars 1 mm in (a,b,e) and 100 µm in (c,d).

**Figure S3. Induction of *pDAD2::GUS* at early stages of lateral root formation.**

**(a)** Very early stage of lateral root primordium with only few visible cells (arrow) and strong local induction of *pDAD2::GUS* in a cortical cell above the primordium. **(b)** Lateral root primordium at a later stage. Root surface is bulging out due to the primordium (arrow) and *pDAD2::GUS* is strongly induced above the primordium. Size bars 50 µm.

**Figure S4. Expression pattern of pDAD2::GUS in roots.**

GUS-stained transgenic roots were fixed, embedded in paraffin, sectioned and processed as described for detailed microscopic analysis. **(a)** Overview of GUS expression in a root with lignified stele. Expression is observed at various sites, with highest levels around the wood in the center. **(b,c)** close up of sites as boxed in (a); arrows mark sites of elevated *pDAD2::GUS* expression. **(d)** Induction of *pDAD2::GUS* in a vascular strand in cells adjacent to xylem vessels evident by their rough appearance (asterisks). Red arrows delimit the vascular strand. Size bars 100 µm in (a), 20 µm in (b,c), 10 µm in (d).

**Figure S5. Responsiveness of petunia roots to exogenous auxin.**

Seedlings were grown on ½ MS plates supplemented with NAA from 1 nm to 1 µM. Primary root growth **(a)** and density of root branching sites (lateral roots and lateral root primordia) (b) were counted after NBT staining. Columns represent the mean (n=6) ± SD (one-way ANOVA and Tukey’s post-hoc test). Columns that share no letter are significantly different.

**Figure S6. Induction of *DR5::3xVENUS-NLS* during early lateral root formation.**

**(a)** Transgenic roots expressing auxin-responsive *DR5::3xVENUS-NLS* exhibit weak nuclear signal throughout the root and elevated levels (bright elongated nuclei; arrows) along the vascular strands in the stele (marked by red autofluorescence). **(b)** High DR5 activity was observed in a cluster of small cells (white arrow) corresponding to a lateral root primordium at a stage corresponding to Fig. 4f. In addition, strong DR5 activity was induced in cells of the root cortex adjacent to the primordium (orange arrows). Red ellipsoid encompasses an area that corresponds roughly to one cell diameter around the root primordium. Size bars 50 µm.

**Figure S7. Induction of *DR5::3xVENUS-NLS* during late lateral root formation.**

**(a)** Transgenic roots show highest *DR5::3xVENUS-NLS* expression in cells around the basis of the LR, reflecting the cells adjacent to the penetration site (green nuclei). **(b)** Image as in (a), but with a tangential orientation along the main root, thereby passing through the base of a lateral root (LR). Cells of the primary root (PR) adjacent to the base of the lateral root (LR) exhibit high levels of *DR5::3xVENUS-NLS* expression. Size bars 50 µm.

**Table S1. Shared induction of genes between auxin and strigolactone (GR24).**

Global gene expression analysis from two independent experiments with hormone-treated petunia roots. A total of 24’816 genes were assessed by microarray analysis as described (Breuillin et al., 2010) and filtered for induction by GR24 (1 µM) of at least 3x above the levels in control roots. Induction ratios (treatment / control) are also shown for treatment with auxin (250 µM IAA) and gibberellic acid (GA_3_) (500 µM).

