## Supplemental Figures for "Strigolactone receptor DAD2 promotes lateral root formation in *Petunia hybrida*"

### Fig. S1. Cell growth adjacent to emerging LRP

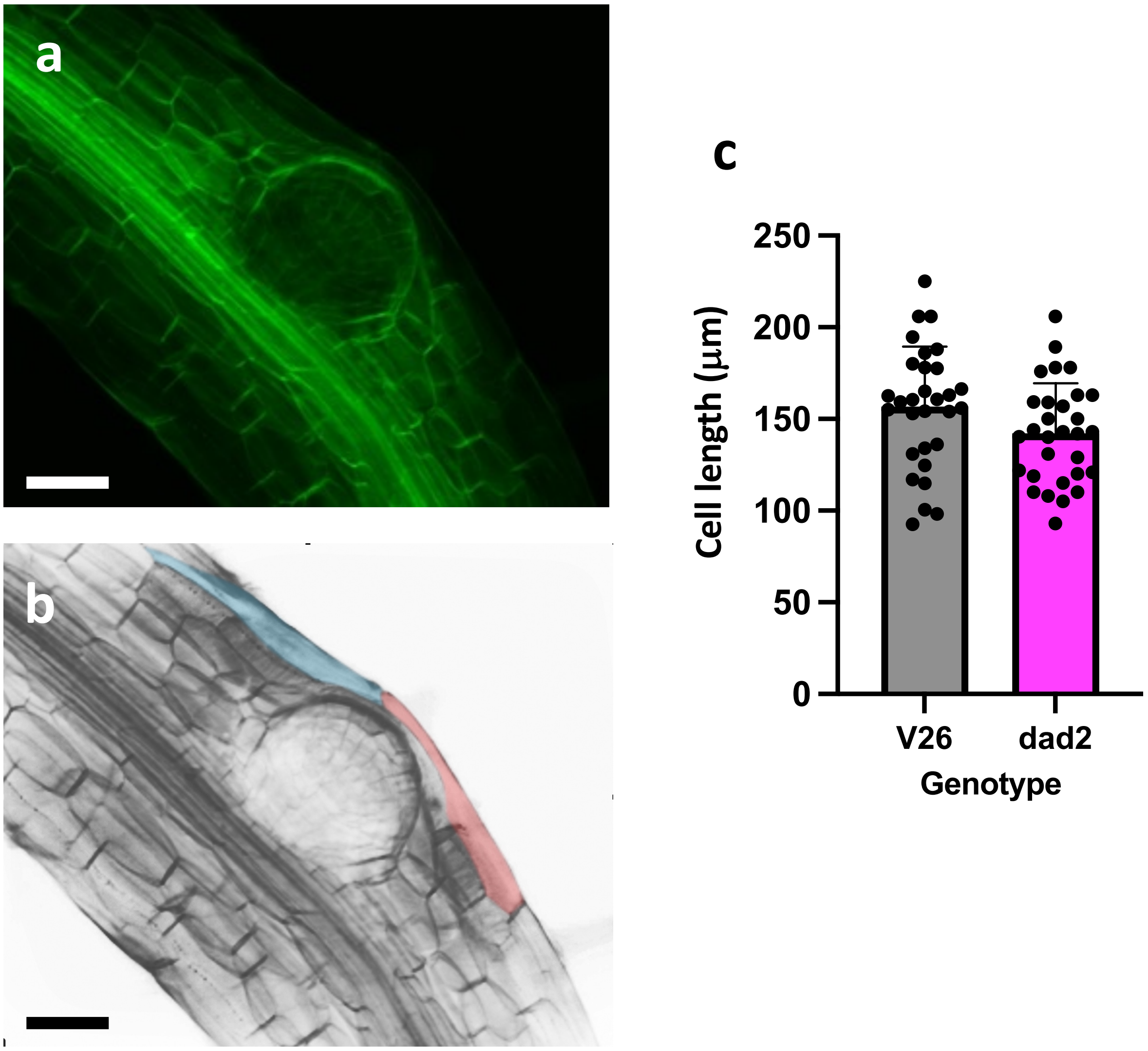

#### Fig. S2. General expression pattern of *pDAD2::GUS*

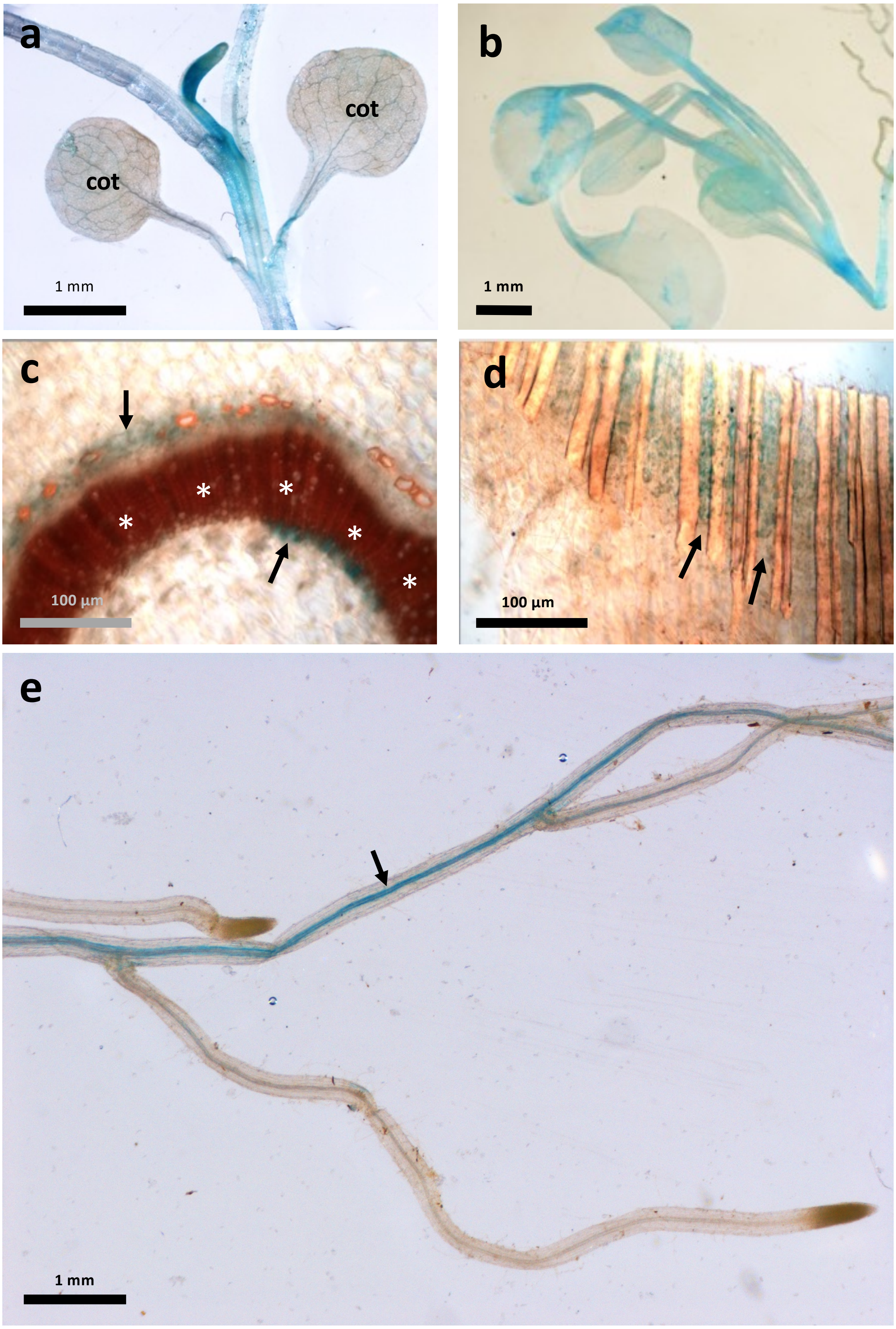

**Fig. S3. Induction of *pDAD2::GUS* at sites or LR emergence**

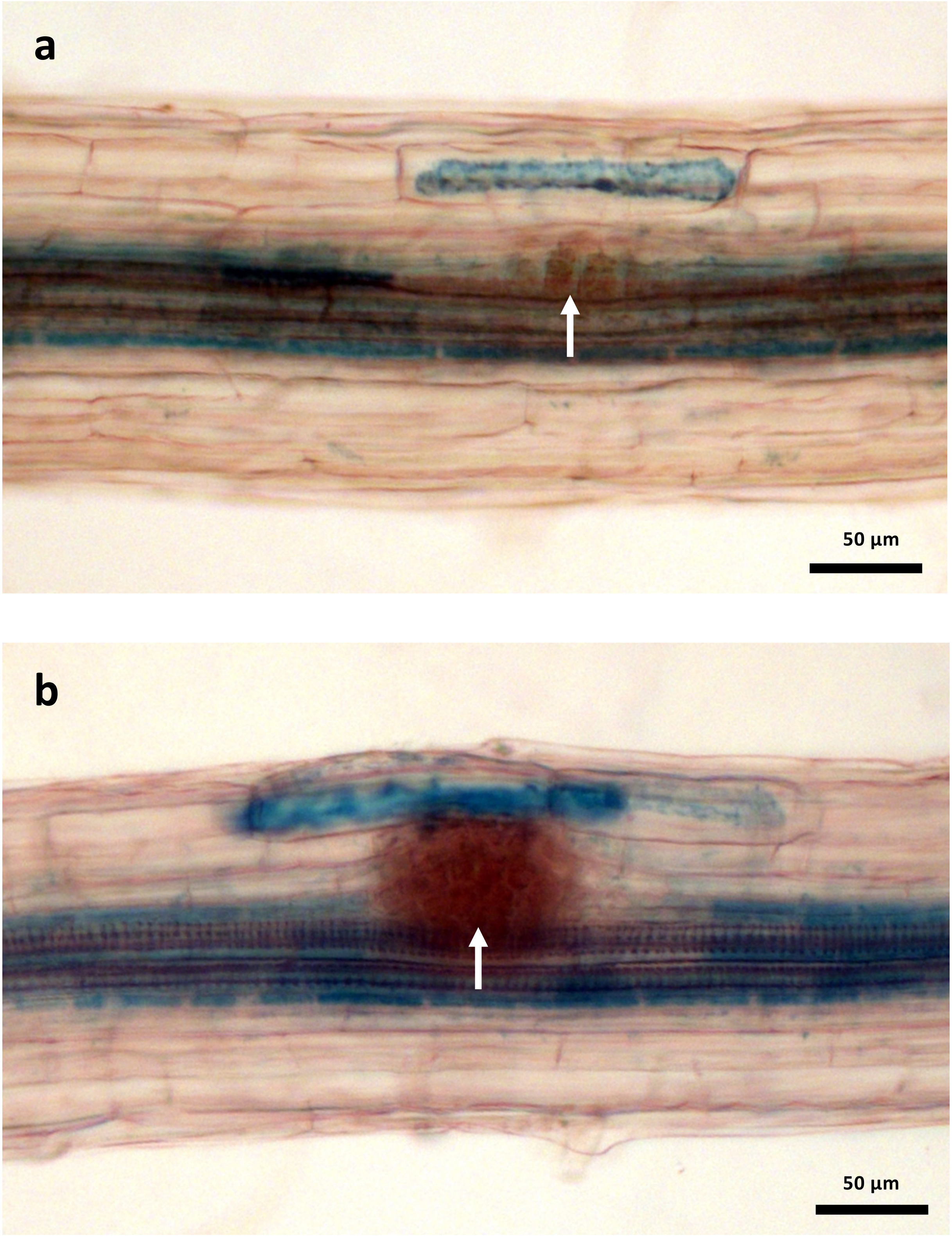

**Fig. S4. Expression of *pDAD2::GUS* in the root**

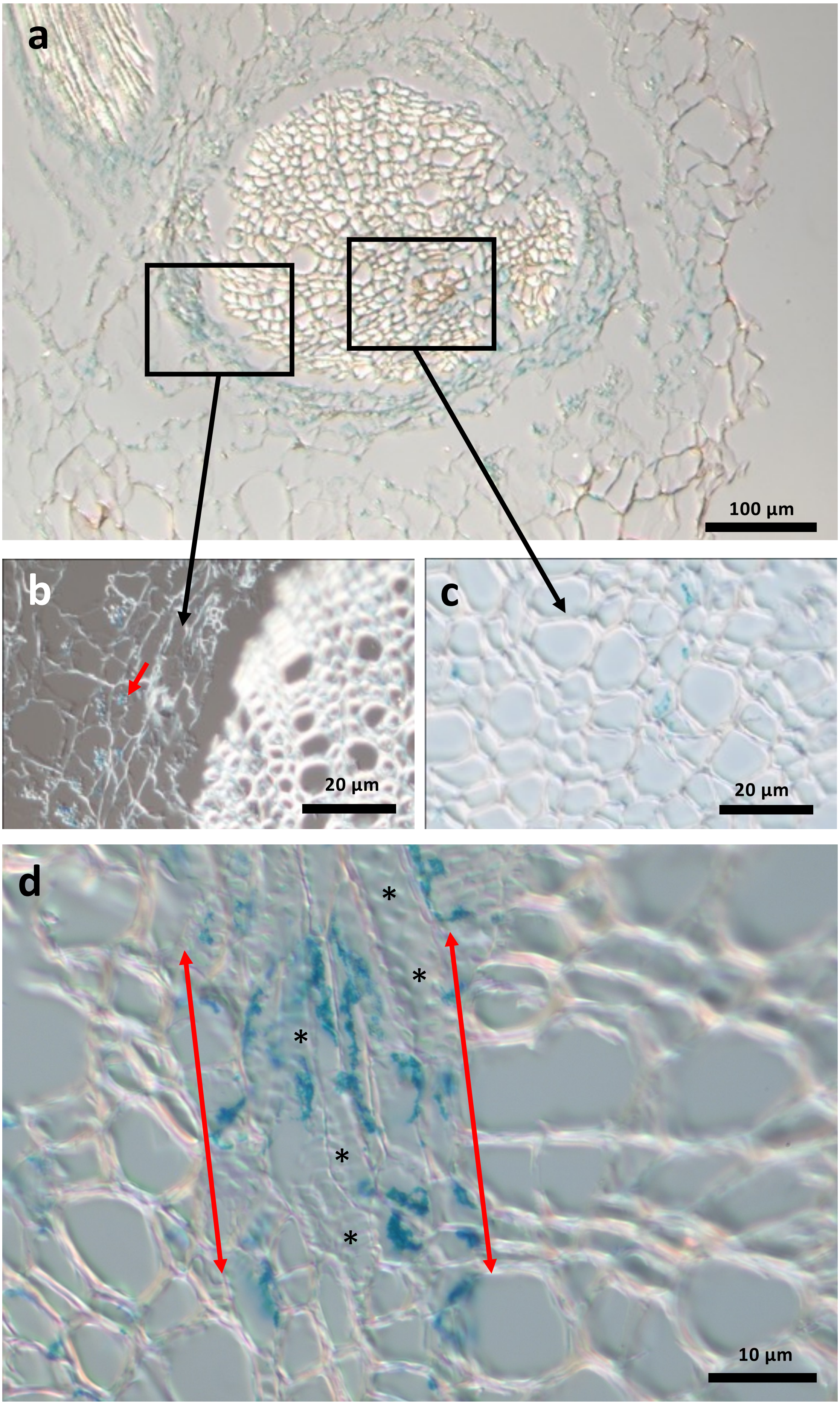

### Fig. S5. Responsiveness of petunia roots to auxin

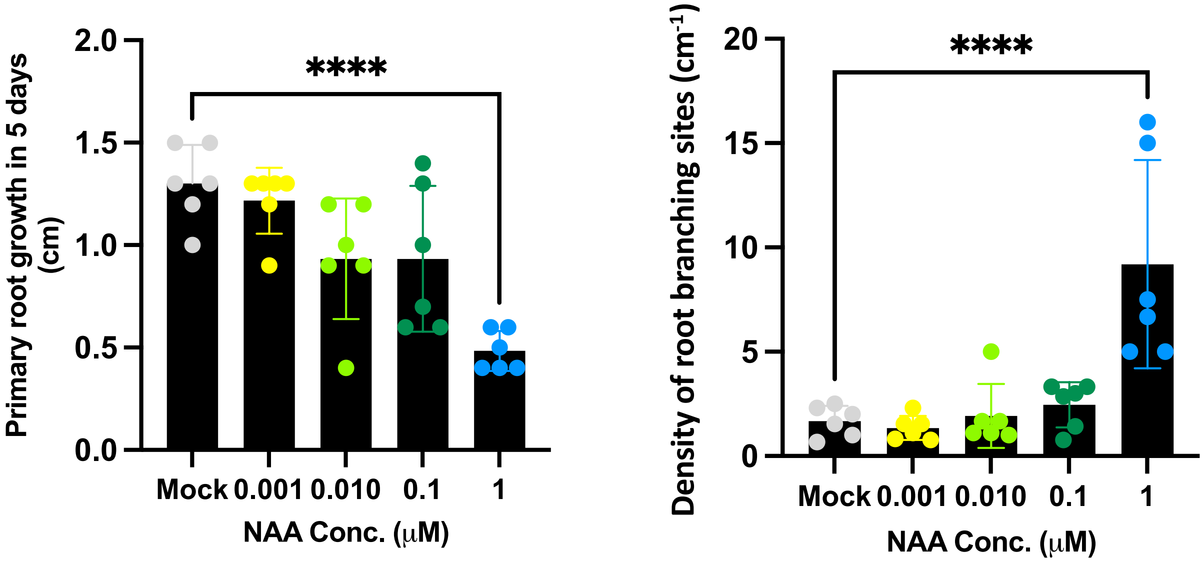

#### Fig. S6. Induction of DR5-NLS-YFP at sites of early LRP

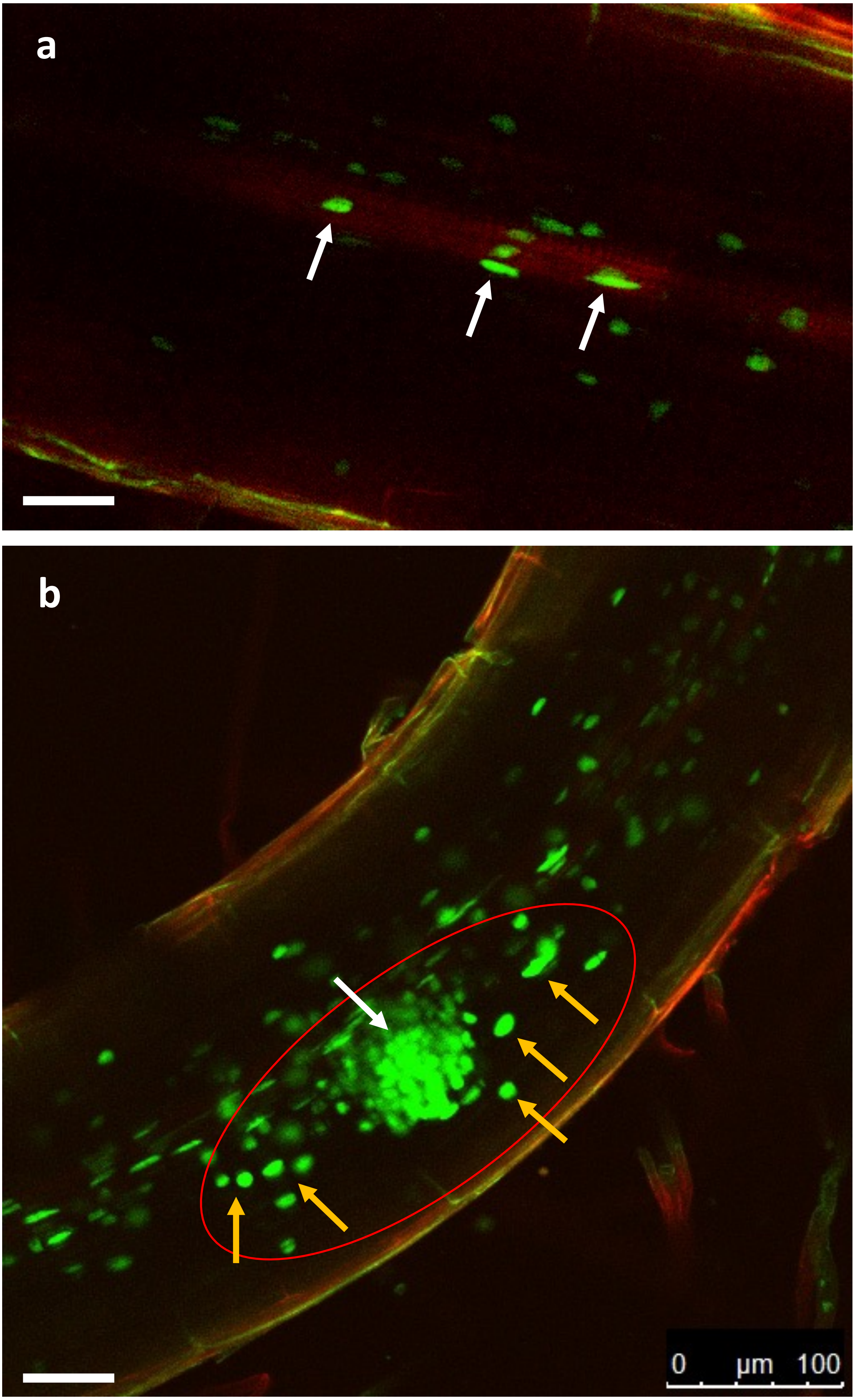

**Fig. S7. Expression of DR5-NLS-YFP at sites of emerged LR**

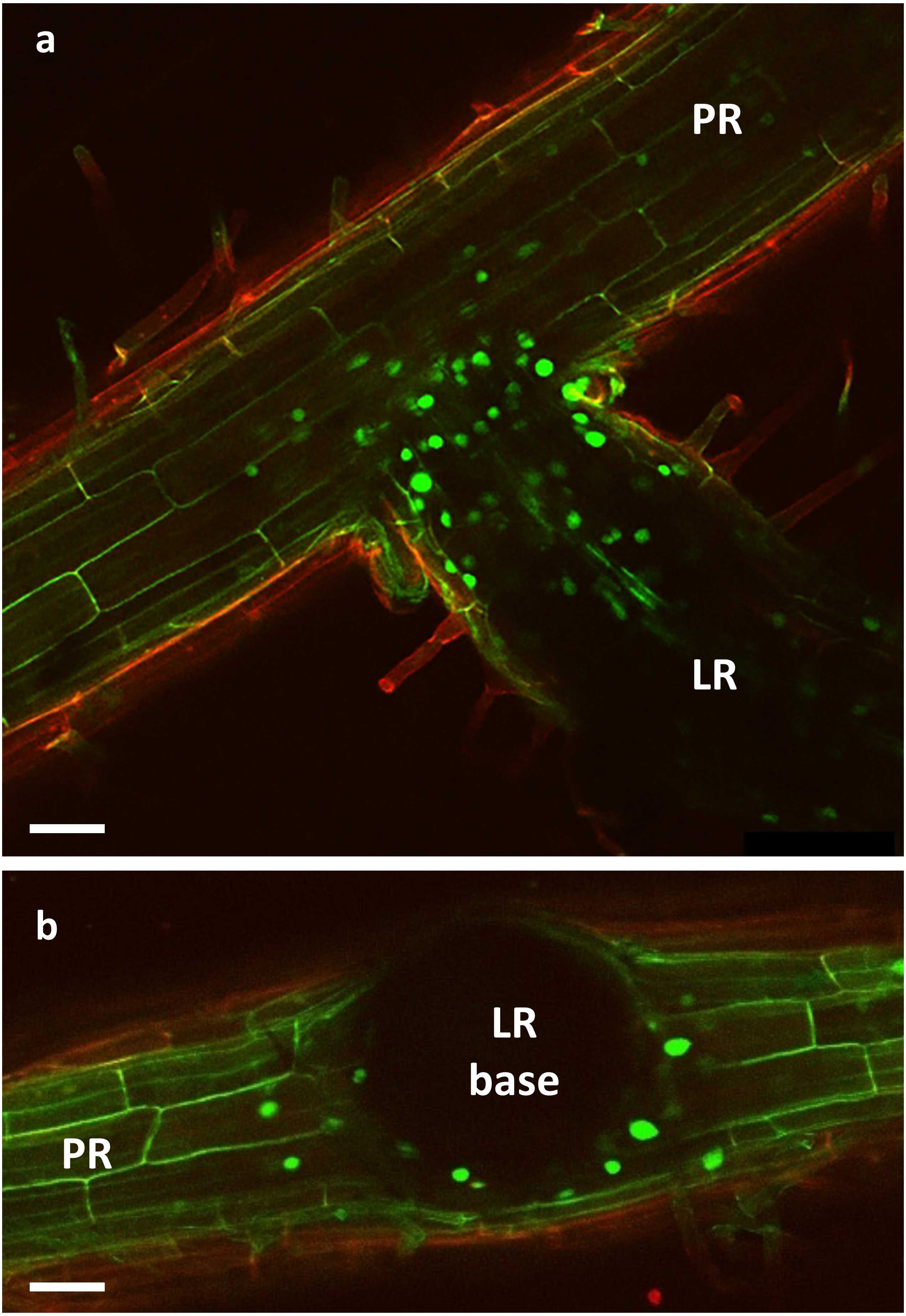

### Table S1. SL-regulated genes in the roots

| Microarray ID | Gene ID in *P. axillaris* (% identity) | Function | Closest homologue in *Arabidopsis thaliana* | *A. thaliana* protein ID | functional class | GR24 1 | GR24 2 | Auxin 1 | Auxin 2 | GA 1 | GA 2 |
|---|---|---|---|---|---|---|---|---|---|---|---|
| GO_dr001P0016M17 | Peaxi162Scf00040g03133.1 (100%) | beta-tubulin | tubulin beta-2/beta-3 chain | BAD93731 | II. Cytoskeleton | 13.39 | 5.59 | 5.02 | 0.77 | 1.39 | 3.12 |
| DY395316_1 | Peaxi162Scf00111g00627.1 (97%) | glutathione S-transferase | GST U26 | OAP16650 | Vb. Antioxidative metabolism and Redox state | 12.85 | 5.82 | 163.04 | 45.70 | 3.01 | 0.48 |
| cn4744 | Peaxi162Scf01082g00111.1 (98.9% | glutathione S-transferase omega | Glutathione-S-transferase family protein | NP_195899.1 | Vb. Antioxidative metabolism and Redox state | 10.79 | 6.31 | 381.06 | 715.87 | 3.63 | 3.25 |
| GO_drs13P0028P05 | Peaxi162Scf00803g00014.1 (88%) | SPX SYG1/Pho81/XPR1 domain-containing protein | SPX domain protein 3 | NP_182038.1 | X. Signalling (Phosphate) | 4.26 | 4.23 | 4.39 | 4.02 | 1.87 | 0.59 |
| cn2706 | Peaxi162Scf00802g00014.1 (95.4%) | fasciclin-like arabinogalactan protein | fasciclin-like arabinogalactan-protein 2 | AAK20858.1 | Ia. Cell wall | 4.20 | 5.72 | 4.72 | 0.71 | 2.26 | 0.79 |
| DC244394_1 | Peaxi162Scf00040g03133.1 (100%) | auxin:hydrogen symporter (PIN-LIKE7) | Auxin efflux carrier family protein | NP_201399.1 | VIb1. Auxin metabolism, transport and perception | 3.92 | 3.10 | 336.40 | 349.30 | 2.90 | 1.70 |
| cn2767 | Peaxi162Scf00104g00083.1 (98.9%) | RuBisCO small subunit | ribulose bisphosphate carboxylase small chain 1A | NP_176880.1 | Va. Primary metabolism | 3.69 | 3.69 | 2.58 | 1.13 | 2.25 | 1.49 |
| cn579 | Peaxi162Scf00016g02023.1 (99%) | caffeoyl-CoA 3-O-methyltransferase | S-adenosyl-L-methionine-dependent methyltransferases superfamily protein | NP_849491.1 | Ia. Cell wall (Lignin) | 3.67 | 5.79 | 15.19 | 2.79 | 4.89 | 0.87 |
| cn8624 | Peaxi162Scf00109g00112.1 (100%) | glutathione S-transferase | glutathione S-transferase tau 7 | NP_180503.1 | Vb. Antioxidative metabolism and Redox state | 3.14 | 4.07 | 28.45 | 38.24 | 1.45 | 1.63 |
